# Chimeric Antigen Receptors Transmit Co-stimulatory Domain Dependent Piconewton Forces to their Target

**DOI:** 10.1101/2025.10.22.683904

**Authors:** Hemakshi J. Mishra, Lauren Mahoney, Dominique R. Smith, Luke Kalcheim, Kartikey Kansal, Nat Murren, Myrna Chang, Rebecca M. Richards, Joshua M. Brockman

**Affiliations:** Biophysics Ph.D. Program, University of Wisconsin—Madison, USA; Comparative Biomedical Sciences Ph.D. program, University of Wisconsin—Madison, USA; Department of Biomedical Engineering, University of Wisconsin—Madison, USA; Department of Pediatrics, University of Wisconsin—Madison, USA; Cellular and Molecular Pathology PhD Program, University of Wisconsin—Madison, USA; University of Wisconsin Carbone Cancer Center – Madison, USA

**Keywords:** DNA Tension probes, Immunotherapy, CAR-T cell therapy, Mechanobiology

## Abstract

Chimeric antigen receptor (CAR) T cells promote tumor-specific cytotoxicity through engagement of a recombinant, synthetic receptor with target ligands expressed on cancer cells. Native T cells are mechanically active, both transmitting and sensing forces exceeding 19 piconewtons (pN) via transmembrane receptors, including the T cell receptor (TCR). Emerging evidence implicates mechanoactivity in CAR T cell biology, but CAR-transmitted T cell forces have not been directly measured. Here, we utilize DNA-based molecular tension probes (MTPs) conjugated to CAR target ligands, providing evidence of actin-polymerization dependent forces exceeding 4.7-19 pN borne by the CAR. We demonstrate force transmission by three clinically relevant CARs (CD123, CD33, and CD19), suggesting that these forces are generalizable across CAR targets and constructs. Additionally, we identify intracellular co-stimulatory domains as the main determinants of CAR-mediated forces, because first-generation CARs lacking co-stimulatory domains do not transmit measurable forces to their ligand. Finally, we demonstrate that CAR forces temporally precede Ca^2+^ signaling and are spatially correlated with phosphorylation of classical TCR-signaling machinery, indicating a link between CAR T cell forces and early biochemical signaling. Our study introduces CAR-mediated mechanobiology as a key correlate of early CAR T cell activation events.

**Significance Statement:** Chimeric antigen receptor (CAR) T cell therapies have revolutionized treatment for several hematological malignancies. CARs are recombinant receptors containing domains derived from the T cell receptor complex machinery and other co-stimulatory proteins. Mechanical forces are believed to be important in T cell activation and antigen recognition. The role of mechanobiology in CAR T cell immunotherapy remains poorly understood. Here, using DNA-based molecular tension probes conjugated to CAR ligands, we provide direct evidence that CARs bear actin polymerization–dependent piconewton forces during antigen engagement that precede early signaling events. These forces depend on CAR co-stimulatory domains, and first-generation CARs lacking these domains fail to transmit detectable force. These findings suggest mechanobiology may be a key, tunable parameter for next-generation CAR T cells.

## Introduction

In chimeric antigen receptor (CAR) T cell therapy, T cells are genetically engineered to express a synthetic receptor that redirects their cytotoxicity towards tumor cells based on interaction with a target ligand^1^. CAR T cell therapies have had striking clinical success for patients with B cell malignancies^2^; however, patient outcomes are varied and translation to myeloid malignancies and solid tumors has been less successful^3^. CAR T cell function is multifactorial and is influenced by many CAR-intrinsic features, including extracellular binding domain affinity, hinge length, and transmembrane and intracellular domain selection^4^. These modular features of the CAR dictate how CAR T cells interact with the tumor and the microenvironment, and influence functional characteristics like anti-tumor efficacy and persistence. Mechanical forces are borne and transmitted by many immune cell receptors, including the T cell receptor (TCR)^5,6^. CARs leverage key features of T cell recognition, but whether mechanical force contributes to CAR signaling is poorly understood.

The earliest “first-generation” CARs comprised extracellular antibody-derived single chain variable fragment (scFv) and hinge regions, linked via a transmembrane domain to the intracellular CD3ζ signaling domain, bypassing major histocompatibility complex (MHC) restriction while still leveraging TCR signaling machinery^7^. In clinical trials, first-generation CAR T cells exhibited poor persistence and tumor control^8^. Subsequently, “second-generation” CARs were designed to incorporate domains from co-stimulatory molecules like CD28 or 4-1BB^3,9^, ultimately improving clinical efficacy and leading to multiple FDA-approved CAR T cell therapies^8^. Generally, CARs bearing a CD28 co-stimulatory domain exhibit faster and more intense activation than their 4-1BB counterparts, which persist longer and retain more memory-like function^10,11^. Exactly how the co-stimulatory domains contribute to CAR T cell short- and long-term function is poorly understood, and CAR development with more complex regulatory signaling and co-stimulatory modules remain areas of active investigation^9,12^.

Native T cells are mechanically active via their TCR. Upon binding to cognate peptide-MHC (pMHC), the TCR actively transmits cytoskeleton-generated forces that regulate T cell signaling and cytokine production^5, 6,12^. The TCR-pMHC forms a “catch bond”, meaning pN forces lead to increased bond lifetime^13,14^. The TCR may associate with actin via residues in CD3ζ^15^, and TCR signaling leads to recruitment of actin polymerization mediators^16^, linking it to cytoskeletal structures. Additionally, the co-stimulatory receptor CD28 may amplify TCR forces through the phosphoinositide 3-kinases (P13K) signaling pathway^17^. A previous study indicates that presenting CAR-ligand on force-cleavable DNA tethers attenuates CAR cytotoxicity and cytokine release^18^, suggesting a role for mechanical force in CAR signaling; however, CAR mechanobiology remains poorly understood.

Given the importance of mechanical forces associated with the TCR, we hypothesized that the CAR likewise transmits forces, and that these forces are regulated by CAR intracellular domains. Among the techniques available to quantify cellular forces^19^, molecular tension probes (MTPs) are particularly well-suited to testing this hypothesis because they allow for direct measurement of the pN-scale forces generated by single receptors as they engage with their target ligand^5, 20^. Accordingly, we employed CAR ligand-conjugated DNA-based MTPs to measure pN-range CAR-mediated forces. We compared forces transmitted by first- and second-generation CARs with the most common co-stimulatory and activation domains and relate these forces to classical early T cell signaling pathways. Here, we provide the first direct evidence that CARs transmit forces to their target antigen through engagement of the T cell cytoskeleton. We also demonstrate that co-stimulatory domains, and not CD3ζ alone, are necessary for CAR-generated forces. Finally, we link CAR mechanics with early biochemical signaling.

## Results

### CAR T Cells Transmit Dynamic, Actin-mediated Mechanical Forces

DNA MTPs comprise a force-extensible DNA hairpin flanked by a fluorophore-quencher pair (**Fig. 1a**). When forces on the MTP exceed the equilibrium force that leads to a 50% chance of hairpin unfolding (F_1/2_), the hairpin opens, separates the fluorophore from the quencher, and produces a “turn-on” increase in signal that is detectable via fluorescence microscopy (**Fig. 1a**). Previously reported DNA MTPs^6,20^ were synthesized, covalently linked to glass substrates, and conjugated with biotinylated CAR ligands via streptavidin (**SI Appendix, Fig. S1**). CD123-specific CAR T cells with a CD28 hinge, transmembrane, co-stimulatory domain and CD3ζ signaling domain (CD123.28ζ) were generated from healthy human donors (**SI Appendix, Fig. S2**), seeded on 4.7pN F_1/2_ DNA MTP substrates presenting CD123, and imaged via total internal reflection fluorescence microscopy (TIRFm). Shortly after contacting target ligand-conjugated MTPs, CAR T cells initiated and maintained contact with the coverslip and generated MTP fluorescence (**Fig. 1b**), indicating that CD123 CARs transmit forces exceeding 4.7pN to their target ligand. As expected, untransduced (Mock) T cells did not transmit any measurable forces to CD123-conjugated MTPs (**Fig. 1c, d**). Additionally, CD123.28ζ CARs did not generate measurable forces on target ligand-mismatched, CD19-conjugated MTPs (**SI Appendix, Fig. S3**). CAR T cell forces were highly dynamic, beginning shortly after cell-surface contact (**Fig. 1e, f**). Most commonly, CAR T cells were stably engaged during the short imaging period, and CAR forces were located at or near the periphery of the cell-surface contact area (**SI Appendix, Fig. S4**). We also observed that several CAR T cells migrated during imaging, and CAR-mediated forces in those cells generally occurred opposite the direction of migration (**Fig. 1e-g**). Collectively, these data demonstrate that CD123 CAR T cells transmit CAR-mediated, spatially heterogeneous, dynamic pN forces to their target ligand.

**Figure 1.**
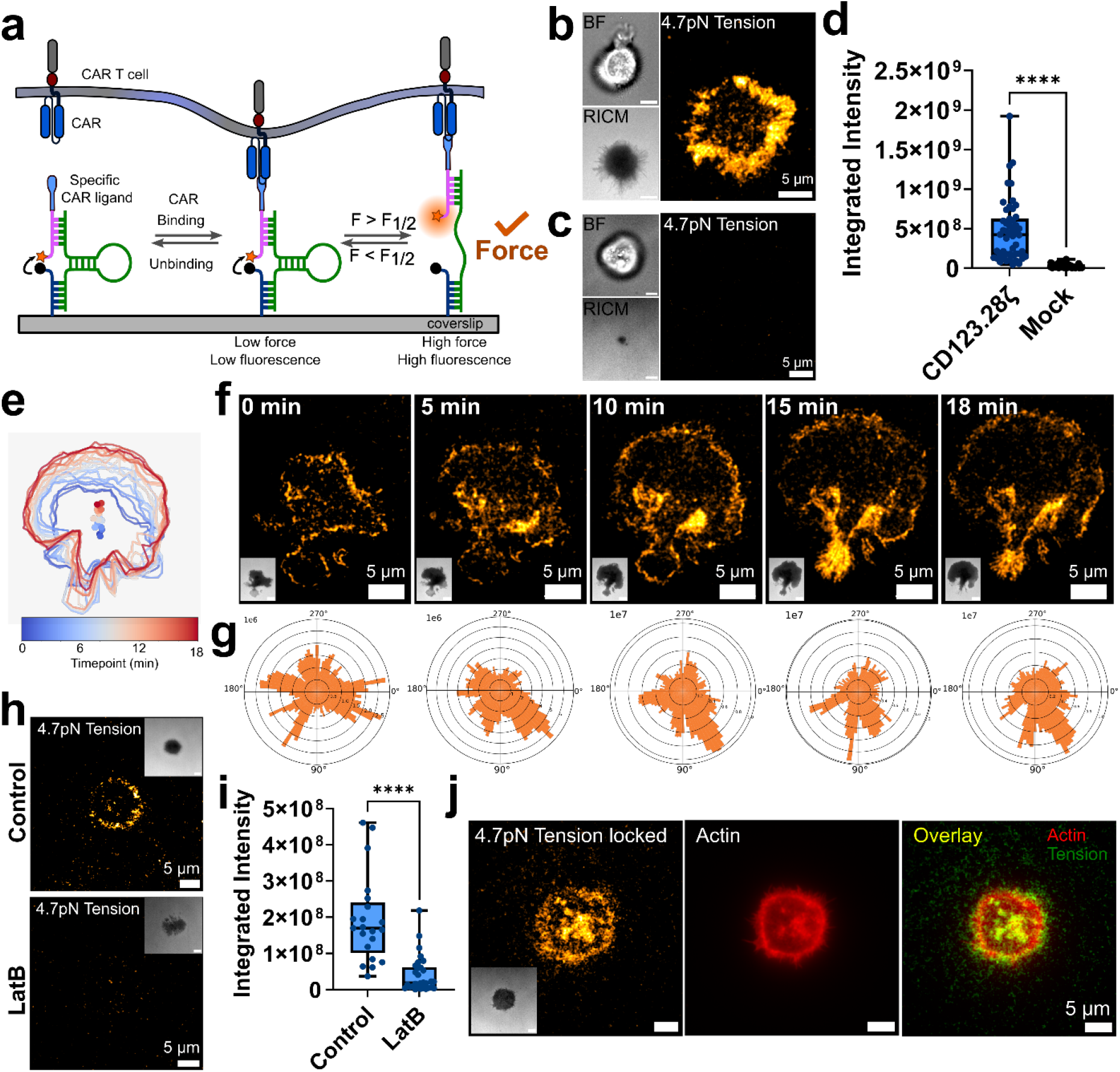
CAR T cells transmit forces to their target ligands. **(a)** Schematic of a DNA MTP presenting a CAR ligand. Forces on the MTP exceeding the probe F_1/2_ rupture the hairpin, producing a turn-on increase in fluorescence. Brightfield (BF), reflection interference contrast microscopy (RICM), and 4.7pN MTP fluorescence image of **(b)** CD123.28ζ CAR T cell and **(c)** mock-transduced T cell seeded on CD123-conjugated MTPs. Images in **b** and **c** are representative of *n* = 3 donors. **(d)** Quantification of whole cell integrated MTP fluorescence for CD123.28ζ CAR (63 cells) and mock T cells (60 cells) seeded on MTPs (n = 3 donors). **(e)** Plot depicting the location of the edges of a CD123.28ζ CAR T cell from 0-18 minutes as it spread on a CD123-conjugated MTP surface. The color of the lines indicates time of observation, while dots indicate the cell’s centroid position as a function of time. **(f)** Timelapse 4.7pN MTP fluorescence images of cell shown in **e**. RICM images are inset. **(g)** Angular histogram of the distribution of MTP fluorescence for the timelapse images in **f. (h)** RICM, and 4.7pN MTP fluorescence images and **(i)** quantification of a CD123.28ζ CAR T cell at t=10 min post addition of LatB or vehicle control. **(j)** 4.7pN MTP fluorescence (locked 1 min, see materials and methods), RICM (inset), actin staining via Alexa647-phalloidin, and red/green overlay with tension map. Images in **j** are representative of *n* = 3 independent experiments conducted with CARs generated from 1 human donor. **** p<0.0001, Mann-Whitney test. The analysis in **e-g** is representative of ∼20 cells from *n* = 3 independent experiments. Scalebars in BF and RICM images in **b**,**c**,**f**,**h** and **j** are 5µm.

To understand how CARs generate mechanical forces, we investigated their cytoskeletal dependence. As TCR-mediated forces are primarily driven by actin polymerization^5^, we asked whether a similar mechanism underlies force generation by CARs. Treating CD123.28ζ CARs with the F-actin polymerization inhibitor LatrunculinB (LatB) abolished MTP fluorescence (**Fig. 1h, i**), suggesting CAR forces depend on actin polymerization. In contrast, as has been observed in endogenous T cells^5^, treatment with inhibitors of myosin contractility did not significantly alter MTP fluorescence (**SI Appendix, Fig. S5**), suggesting that CAR forces are not dependent on actomyosin contractility. Additionally, fixed MTP fluorescence colocalizes with the actin cytoskeleton, further supporting the importance of actin in CAR force generation (**Fig. 1j**).

### Multiple CAR-Target Ligand Pairs Generate pN Forces

We next sought to extend these findings beyond CD123 CAR T cells to determine whether pN forces are generalizable across a variety of clinically relevant CAR-ligand pairs. Accordingly, we generated CAR T cells specific for CD33 or CD19, both including CD28 co-stimulatory domains (CD33.28ζ or CD19.28ζ, respectively). CD33, like CD123, is a common acute myeloid leukemia CAR target^21^, and CD19 is the most widely studied CAR target given its uniform and high expression in many B cell malignancies^1^. To increase the sensitivity of CAR T cell force measurement, we employed a previously published^22^ signal amplification strategy in which a complement to the DNA hairpin of the MTP binds to mechanically opened hairpins, “locking” them in the open conformation (**Fig. 2a**). This locking strategy integrates the total mechanical output of cells over time, amplifying MTP fluorescence and increasing the sensitivity of MTPs to transient or rare force events. We seeded CD123.28ζ (**Fig. 2b**), CD33.28ζ (**Fig. 2c**), and CD19.28ζ (**Fig. 2d**) CAR T cells onto surfaces conjugated with DNA MTPs bearing the corresponding target ligand. 4.7pN F_1/2_ MTP fluorescence signal was readily apparent for CD123.28ζ (left, **Fig. 2b**) and CD19.28ζ (left, **Fig. 2d**) CARs in real time. In contrast, CD33.28ζ CARs generated significantly weaker real time MTP fluorescence (left, **Fig. 2c**). The locking strategy amplified the MTP signal generated by all CARs and improved our ability to visualize forces transmitted by the CD33.28ζ CAR (**Fig. 2c**). We quantified the integrated per-cell MTP fluorescence signal for real time (**Fig. 2e**) and locked MTP signal (**Fig. 2f**), revealing that CD33.28ζ CARs transmit fewer forces exceeding 4.7pN via their chimeric receptors than CD19.28ζ and CD123.28ζ CAR T cells. Interestingly, CD123.28ζ CAR force generation is more abundant than the gold-standard CD19.28ζ CAR in real time MTP measurements (**Fig. 2e)** and locked MTP measurements (**Fig. 2f**). These results suggest that mechanical forces may be a common feature across CAR T cells but also suggest that CAR T cell forces may vary by scFv-target ligand pairing.

**Figure 2.**
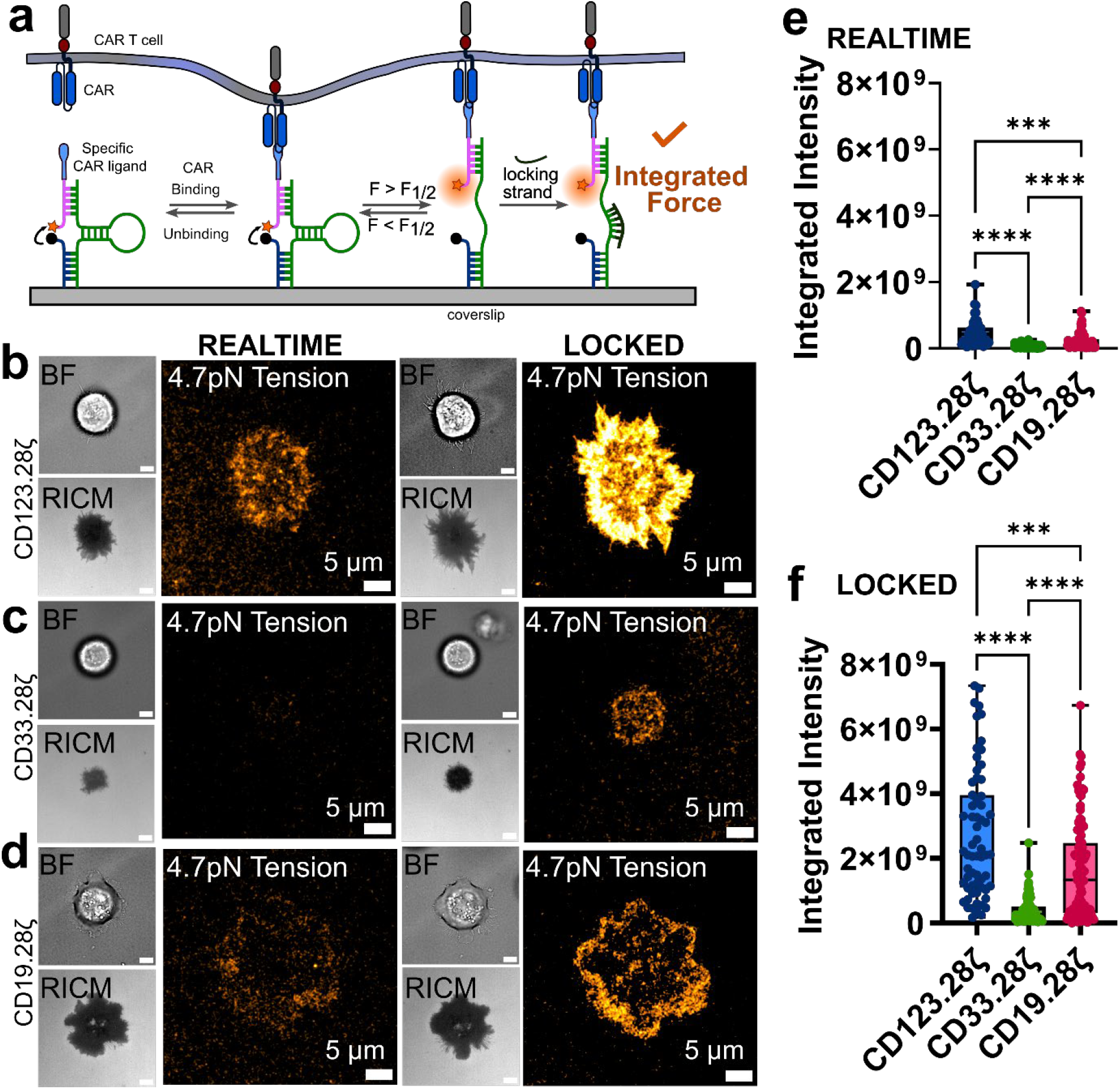
Forces exerted by CARs vary by CAR-target ligand pairing. **(a)** Schematic of locking strategy that integrates force transmission events. BF, RICM, and 4.7pN MTP fluorescence images of **(b)** CD123.28ζ, **(c)** CD33.28ζ and **(d)** CD19.28ζ CAR T cells seeded on their respective target ligand-presenting MTPs. Quantification of **(e)** realtime and **(f)** locked MTP fluorescence for CD123.28ζ, CD19.28ζ, CD33.28ζ CAR T cells from *n = 3* human donors. Images in **b, c** and **d** are representative of *n* = 3 human donors. *** p<0.001 **** p<0.0001, Kruskal-Wallis test. Scalebars in BF and RICM images in **b-d** are 5µm. Note that CD123.28ζ quantification data from **Fig. 1d** is replotted in **2e** for ease of comparison with CD33.28ζ and CD19.28ζ data.

### Co-stimulatory Domains Drive Force Generation

All FDA-approved CAR T cells incorporate either a CD28- or a 4-1BB-derived co-stimulatory endodomain, along with the intracellular domain of CD3ζ, which propagates signaling through its immunoreceptor tyrosine-based activation motifs (ITAMs)^23^. Although the CAR is mechanically active, it is not clear whether the CD3ζ domains incorporated into the CAR are sufficient to generate CAR-mediated forces, or whether the co-stimulatory domains drive or augment force. Accordingly, we sought to assess the role of CAR-intrinsic co-stimulatory domains on mechanical force generation. Both CD123.28ζ (**Fig. 3a**) and CD123.BBζ (**Fig. 3b**) CARs produced MTP tension signal. Interestingly, first-generation CD123 CAR T cells, which incorporate a CD28 hinge/transmembrane domain and CD3ζ signaling domain but have no co-stimulatory domain (denoted CD123ζ), failed to generate MTP fluorescence, whether measured in real-time measurements or with the locking strategy (**Fig. 3c**). Quantification of integrated MTP fluorescence suggests CD123.28ζ CAR T cells are more mechanically active than CD123.BBζ cells, while both CD123.28ζ and CD123.BBζ CARs transmit more forces than first-generation CD123ζ CAR T cells (**Fig. 3d**). In fact, the MTP fluorescence of CD123ζ CARs matched that of mock T cells (**SI Appendix, Fig. S6**), This failure to generate force was not due to low CAR expression levels (**SI Appendix Fig. S6**), and CD123ζ CARs maintained equivalent cytotoxicity to second generation CARs *in vitro* (**SI Appendix, Fig. S6**). Similarly, CD19.28ζ CARs transmitted more MTP fluorescence than CD19.BBζ CARs (**SI Appendix, Fig. S7**). No statistical difference was observed between the CD33.28ζ and CD33.BBζ CARs (**SI Appendix, Fig. S7**), possibly due to due to their low overall MTP fluorescence. Additionally, both CD123.28ζ and CD123.BBζ CARs produced MTP tension signal on probes with F_1/2_= 19pN (**SI Appendix, Fig. S8**). These differences did not correlate with variation in CAR transduction (**Fig. 3e**), or the CD4/8 ratio (**SI Appendix, Fig. S9**). Collectively, these findings implicate CAR co-stimulatory domains, and not CD3ζ, as key regulators of CAR-mediated forces.

**Figure 3.**
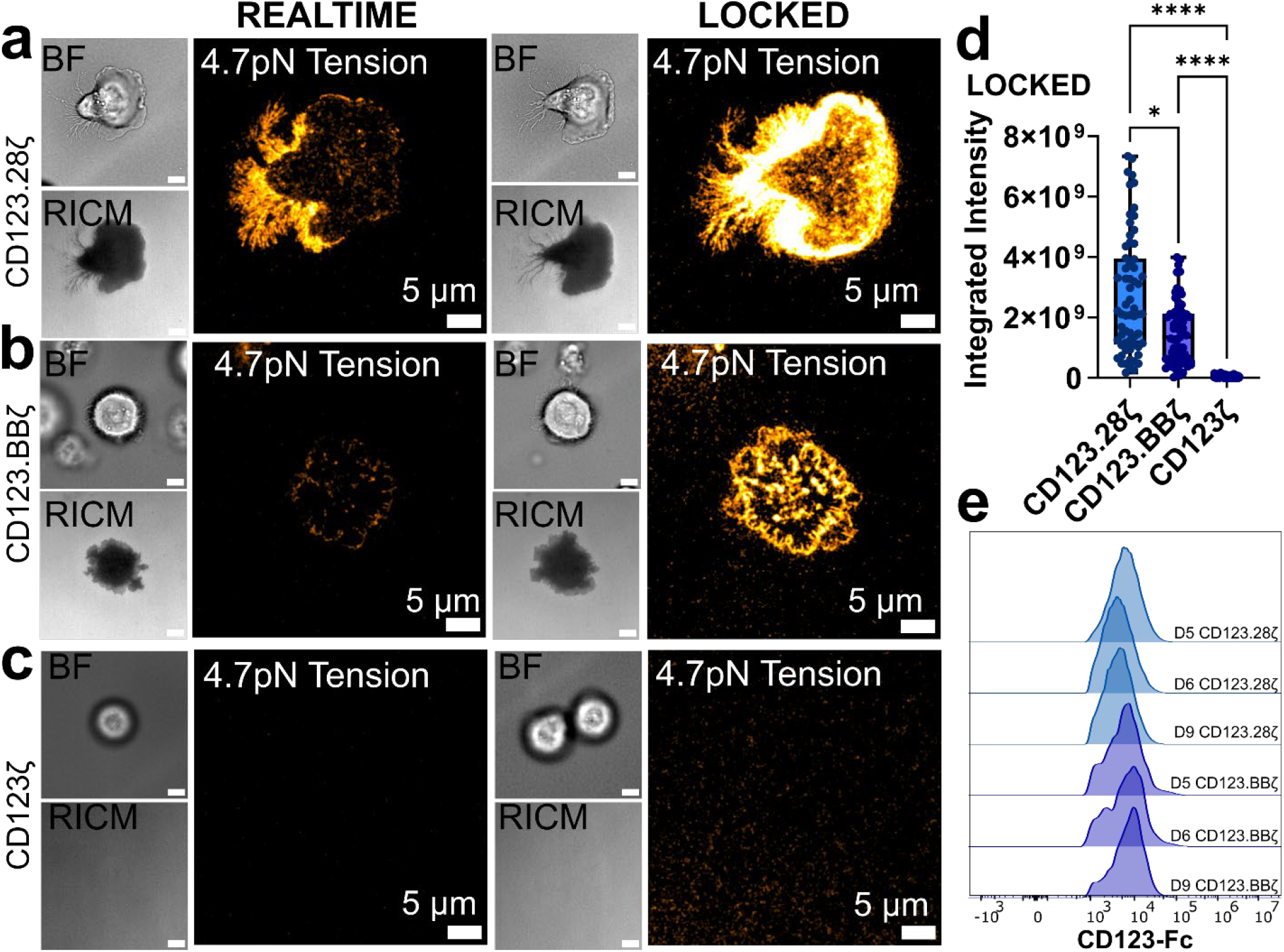
CAR forces depend on the co-stimulatory domain. BF, RICM, and 4.7pN MTP fluorescence image of **(a)** CD123.28ζ, **(b)** CD123.BBζ, and **(c)** CD123ζ CAR T cells seeded on CD123-presenting MTPs. **(d)** Quantification of MTP fluorescence for all CD123 CARs from *n* = 3 human donors. **(e)** Intensity of CD123 CAR expression measured via flow cytometry. Images in **a-c** are representative of *n* = 3 human donors. * p<0.05, **** p<0.0001, Kruskal-Wallis test. Scalebars in BF and RICM insets in **a-d** are 5µm. Note that CD123.28ζ locked quantification data from **Fig. 2f** is replotted in **3d** for ease of comparison with CD123.BBζ and CD123ζ data.

We also quantified the integrated MTP fluorescence signal of CAR T cells from multiple de-identified human donors (**SI Appendix, Fig. S10**). We found that CAR MTP tension signals vary among donors. Additionally, while not universally true, we observed that across many constructs, donor 9 CAR T cells were the most mechanically active. (**SI Appendix, Fig. S10**). Currently, the origin of donor-to-donor variability in MTP signal remains unclear; however, it is possible that genetic and environmental factors that influence clinical CAR T cell activity^24^ may also influence CAR-mediated forces.

### Mechanical Forces Precede and Correlate with Early T Cell Signaling

Next, we sought to determine whether CAR-mediated forces correlate with a polyfunctional, pro-inflammatory cytokine signature. We performed intracellular cytokine staining for IFNγ, TNFα, perforin, and granzyme B after CD123, CD33, and CD19 CAR T cells from multiple donors were stimulated with beads presenting their corresponding ligands (**SI Appendix, Fig. S2, S11**). The average MTP fluorescence of all measured cells from a given CAR-donor pairing correlates with the capacity of that CAR T cell product to produce IFNγ and TNFα upon antigen stimulation. (**SI Appendix Fig. S11**). We did not observe a significant correlation between MTP fluorescence and granzyme B or perforin (**SI Appendix, Fig. S11**). First generation CAR T cells, which fail to generate MTP fluorescence, do not exhibit the polyfunctional cytokine signature that is seen in second generation CAR T cells (**SI Appendix, Fig. S11f-h**).

To measure force and its effect on immediate downstream T cell signaling events in our MTP, single cell system, we imaged early biochemical markers of T cell activation. Calcium (Ca^2+^) influx is an early process in T cell activation^25^. Accordingly, we stained CD123.28ζ CARs with the calcium sensitive dye Fluo-4 AM and subjected them to timelapse imaging (**Fig. 4a**). MTP fluorescence was observed shortly after CARs made surface contact; however, Ca^2+^ influx was only observed after MTP fluorescence (**Fig. 4a-c**). We measured the mean MTP fluorescence intensity (black trace, **Fig. 4b**) and integrated Fluo-4AM intensity (pink trace, **Fig. 4b**) for 27 CAR T cells from *n*=3 independent experiments. We defined a parameter, Δt_rise_, as the time delay between MTP fluorescence onset and Ca^2+^ influx (Δt_rise_ = 90s for the cell shown, **Fig. 4a, b**). We measured an average Δt_rise_ of 175 seconds (**Fig. 4c**). Importantly, Ca^2+^ influx was never observed before MTP fluorescence. To further assess early biochemical signaling events, we fixed and stained CD123.28ζ CARs for phosphorylated Zap70 (pZap70). Zap70 is recruited and phosphorylated during initial T cell signaling upon antigen recognition^26^. pZap70 spatially correlates with MTP fluorescence (**Fig. 4d, e**) and per cell integrated tension fluorescence intensity is positively correlated with pZap70 staining (**Fig. 4f**). These data indicate that CAR forces precede, and are associated with, early T cell signaling.

**Figure 4.**
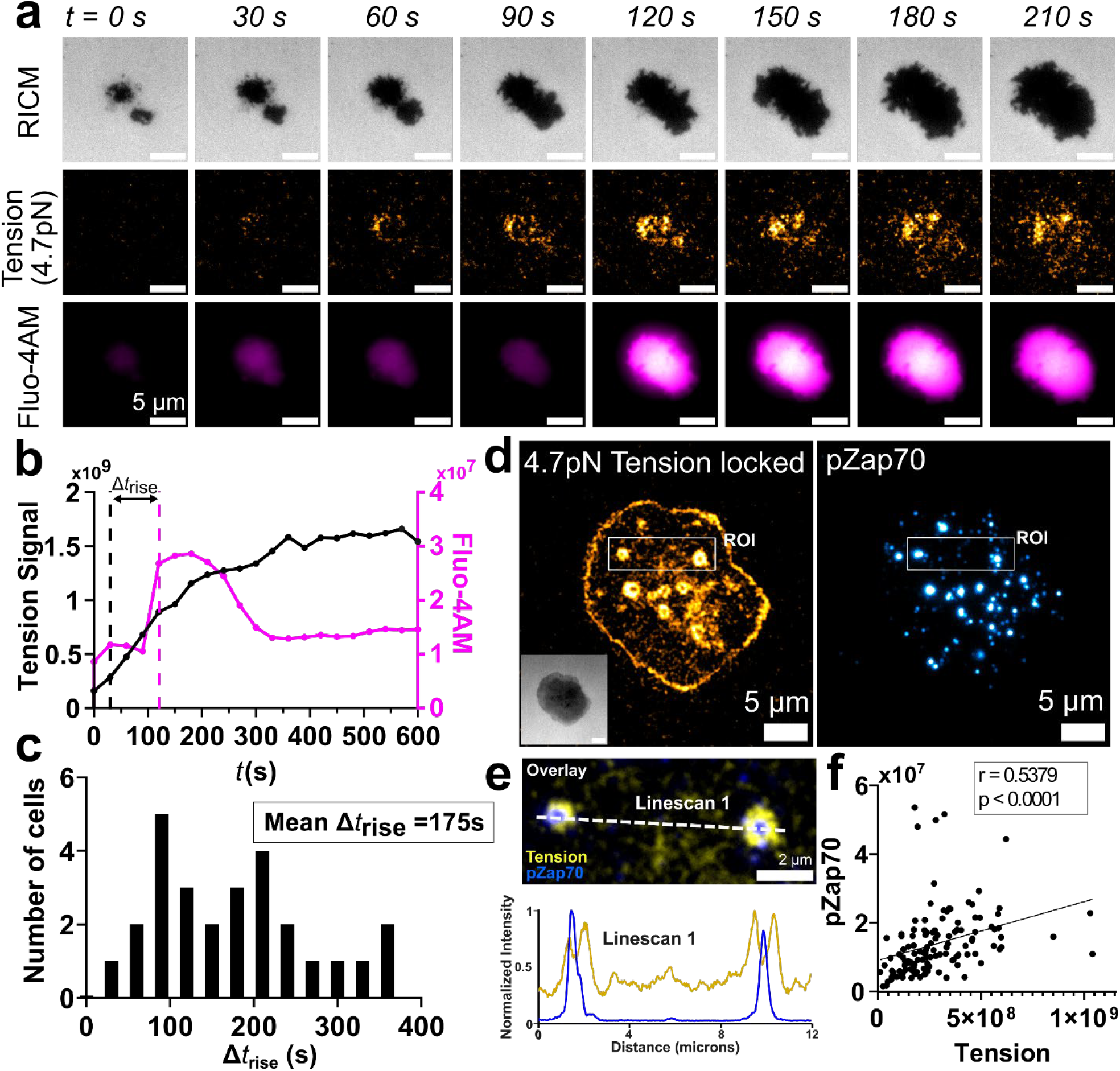
CAR-mediated forces correlate with early CAR T cell signaling events. **(a)** Representative timelapse images and **(b)** whole cell intensity trace of tension signal and fluo-4 AM fluorescence showing temporal differences between the initial rise of [Ca2+] and the initial rise in CAR tension (Δt_rise_) for a CD123.28ζ CAR T cell. **(c)** Histogram analysis of Δt_rise_ (n = 27 cells). **(d)** RICM (inset), 4.7pN MTP fluorescence (locked) and pZap70 fluorescence image of a CD123.28ζ CAR T cell on CD123 presenting MTPs. **(e)** Zoom in image of ROI indicated in **d**. Intensity linescan along the dotted white line is provided. (**f**) Correlation of per cell integrated tension signal with integrated pZap70 fluorescence intensity. Data in **a-f** are representative of *n* = 3 independent experiments. Scalebars in RICM images in **a** and **d** are 5µm.

## Discussion

Our findings establish that, like native TCRs, synthetic CARs are mechanically active. Although prior studies have hinted at CAR mechanosensitivity^18^, we provide direct MTP-based evidence of force transmission through the CAR, raising the possibility that upon CAR-target ligand engagement, downstream signaling is mechanically regulated. The mechanosensitivity of T cells via their TCR is well-appreciated. Native T cells transmit forces exceeding 19pN via their TCR^5^. Their activation depends on the stiffness^27,28^ and viscoelasticity^29^ of the stimulating substrate. Additionally, T cell cytotoxicity is increased against cancer cell seed on stiff substrates^12^, while biologically soft cancer cells impede T cell cytotoxic activity^30^. Our data support the mechanobiology of CAR–target ligand interactions as a novel factor that may influence CAR T cell function. Indeed, we provide correlative evidence of CAR mechanics with early signaling events. We show that CAR forces precede calcium signaling, and that those forces spatially correlate with pZap70, a critical kinase in T cell signaling. Given these findings, further study of CAR mechanotransduction is warranted.

Co-stimulatory domains modulate CAR T cell activation, signaling, and functional potency^10,11,31^. We present data that co-stimulatory domains also regulate CAR-mediated forces. Both CD123.28ζ and CD123.BBζ CARs transmit forces exceeding 19pN to their ligand. Therefore, at the protein level, CARs with 4-1BB co-stimulatory domains are not mechanically weaker than those with CD28 co-stimulatory domains, at least within the bounds of our MTP force sensitivity. Our data, which show higher MTP fluorescence signal for CD28 than 4-1BB CARs, are consistent with several possible explanations: (1) a greater fraction of CD28ζ-containing CARs are mechanically active; (2) 4-1BBζ-containing CARs generate transient forces, analogous to a low “duty cycle” of mechanical activity, limiting detection; (3) 4-1BBζ-containing CARs sample fewer MTPs over time; or (4) a combination of these mechanisms.

Interestingly, first-generation CARs—lacking any co-stimulatory domain—do not generate measurable forces. The inability to transmit forces above 4.7pN may be one factor that contributes to their limited clinical efficacy^32^. In the context of CARs, CD28 co-stimulatory domains are known to have faster interaction times and are generally more rapid in their cytotoxicity and expansion after target engagement^10^. Previous experiments have shown that Cd28 augments TCR-based forces^17^. Additionally, CD28 engagement promotes actin polymerization via Cdc42^33^, providing a potential direct link to the actin cytoskeleton. In contrast, 4-1BB as a co-stimulatory has not been linked to T cell force to the best of our knowledge^34^, making our measurement of 4-1BB CAR forces fertile ground for future inquiry.

Our data additionally reveal that sites of mechanical tension within CAR T cells spatially coincide with actin-rich foci, and that disrupting actin polymerization with LatB abolishes these forces, implicating actin dynamics as the primary driver of CAR-generated tension. These tension-rich foci may represent mechanically active CAR assemblies, as we detect activation-associated pZap70 hubs at these same sites. This aligns with previous studies showing that CAR immunological synapses are multifocal and less organized than the standard concentric structure of cytotoxic TCR immunological synapses^35, 36^. Notably, conventional T cells form F-actin–rich protrusions at the synapse that are well-recognized as mediators of force transmission, signaling and cytotoxicity^37–39^. Together, these observations support that CARs, likewise, form actin-rich, mechanically active structures at the synapse that may promote rapid proximal signaling. Our findings also suggest a correlation between CAR mechanical forces and CAR T cell production of IFNγ and TNFα. A link to cytotoxicity is less clear. For example, first-generation CAR T cells fail to generate measurable forces but do kill target cells *in vitro*, indicating that lack of mechanical force through the CAR is not a complete limitation for CAR T cell cytotoxicity in reductionist systems. It is possible that in complex, mechanically diverse milieus such as the solid tumor microenvironment, CAR-mediated forces may be crucial. Further investigation of a link between CAR forces and target lysis in different contexts is warranted.

The consequences of CAR-transmitted forces remain unclear. While CARs integrate TCR-derived CD3ζ as a signaling module, they are fundamentally distinct immunoreceptors. The TCR–pMHC forms a catch bond^13,14^, allowing force to stabilize antigen engagement^40^ and promote structural transitions such as in the TCR FG-loop region^41^. This behavior enables the TCR to use mechanical load as part of its antigen discrimination process. By contrast, most antibody–antigen (scFv–ligand) interactions form slip bonds^42^ whose lifetime decreases under force. For most CAR scFvs, slip bonds likely cause force application to accelerate bond dissociation. Mechanically, this sets up an interesting paradox: CARs use intracellular machinery (e.g., CD3ζ ITAM phosphorylation, actin coupling) that can generate and respond to TCR-like pN-scale forces, but the extracellular interface is not designed to bear those loads productively. It remains unclear whether CAR forces ultimately promote or inhibit signaling, and both possibilities have consequences for engineering CAR T cells that optimize bond length and downstream functional outcomes.

Unpredictable bench to bedside translation and interpatient variability remain significant hurdles in CAR T cell therapy^9^. Our findings establish mechanical force as a novel axis that correlates with early signaling events known to be central to CAR T cell function, suggesting that force generation and mechanosensing could be incorporated into the iterative process of preclinical CAR T cell evaluation. By defining how co-stimulatory domains, actin coupling, bond dynamics, and receptor conformations integrate mechanical signals, it may be possible to guide CAR engineering toward designs that optimize signaling and cytotoxicity in clinically relevant mechanical environments. Ultimately, this work highlights the underappreciated role of mechanics in CAR T cell biology and opens new avenues for engineering next-generation immunotherapies that harness both biochemical and biomechanical principles.

## Supporting information

Combined Supplemental Appendix

## Acknowledgements

JMB wishes to acknowledge R35GM157007 and support from ACS grant IRG-19-146-54. RMR acknowledges K08 CA266930 and an ASH Scholar Award.

## Author contributions

HJM, LM, LK, and NM performed experiments. HJM, LM, DRS, MU, KK, and MC performed data analysis and generated figures. JMB and RMR designed and oversaw the study. All authors wrote and edited the paper.

