## Supplementary material for "Chimeric Antigen Receptors Transmit Co-stimulatory Domain Dependent Piconewton Forces to their Target": Combined Supplemental Appendix

**This PDF file includes:**

Materials and methods
Figure S1: Surface preparation scheme
Figure S2: Flow cytometry gating strategy for CAR T cell phenotyping and intracellular cytokine staining
Figure S3: CARs do not transmit force to irrelevant ligand
Figure S4: CARs transmit dynamic forces on MTPs
Figure S5: Forces exerted by CARs are not myosin-mediated
Figure S6: “First-generation” CD123.ζ CARs fail to generate forces exceeding 4.7pN but retain cytotoxicity
Figure S7: Both CD28 and 4-1BB co-stimulatory domain CAR T cells targeting CD123, CD33, or CD19 transmit pN forces to target ligand
Figure S8: CD123 CAR T cell exert forces exceeding 19pN
Figure S9: MTP tension fluorescence comparison with CD4/8 ratios of CAR T cells
Figure S10: Tension of CARs is donor T cell-dependent
Figure S11: Intracellular cytokine staining following CAR target ligand-conjugated bead stimulation
Figure S12. Image processing scheme
Figure S13. Dye labelled DNA strand synthesis
Table S1. List of oligonucleotides
Table S2. List of fluorophores and antibodies utilized in flow cytometry

### Materials and Methods

Materials:

Chemically modified oligonucleotides were synthesized by Integrated DNA Technologies (IDT). Sequences are available in **Supplementary Table 1.**

Cy3b-NHS was acquired from BroadPharm (BP-28886). Phosphate-buffered saline (PBS), acetonitrile, and triethylamine were obtained from Thermo Fisher Scientific. No. 1.5H coverslips (#10812) and sticky-Slide VI 0.4 (#80608) were purchased from Ibidi (Fitchburg, WI). Streptavidin (#50-105-9142) was purchased from Rockland Immunochemicals. Additional reagents, including bovine serum albumin (BSA) and cytoskeletal inhibitors (Latrunculin B #42-802-01 and Y27632 dihydrochloride #Y0503) were purchased from Sigma Aldrich. All buffers were made with Nanopure water (18.2 MΩ) and passed through a 0.2 µm filtration system. Azide-PEG$_4$-NHS ester (#AZ-103) was obtained from Click Chemistry Tools (Scottsdale, AZ). Biotinylated CAR ligands CD123 (ILA-H8214), CD33 (CD3-H82E7) and CD19(CD9-H82F6) were purchased from AcroBIOSYSTEMS. Alexa Fluor 647 Mouse Anti-ZAP70 (PY319)/Syk (PY352) (#557817) was obtained from BD Biosciences and Alexa Fluor Plus 647 Actin Phalloidin stain (#A30107) was obtained from Fisher Scientific.

#### Dye labelled DNA strand synthesis

Probes were synthesized using a modification of previously reported methods[6]. The ligand strand was modified at its 3'-terminus by coupling Cy3B–NHS to the available amine group. Synthesized probes were purified by high-performance liquid chromatography (HPLC) and characterized via mass spectrometry. Representative HPLC and mass spectrometry data are provided in **SI Appendix Fig. S13**.

#### CAR construct production

The heavy and light chain sequences antibody sequences targeting CD123 (32716, patent #WO2014130635A1), CD19 (FMC63, patent #US9701758B2), and CD33 (Hu195, patent #WO2020223445A1) were acquired from the patent data base, synthesized by Integrated DNA Technologies (IDT, Coralville, IA) and linked with a $(G_4S)_3$ sequence. Standard restriction enzyme digestion and ligation techniques were used to insert the ScFV sequences into a MSGV1 backbone containing either a CD8a derived hinge and transmembrane sequence with a 4-1BB derived intracellular portion or a CD28 derived hinge, transmembrane, and intracellular domain, each with a distal CD3ζ intracellular signaling domain. The CD123ζ first-generation CAR was produced with similar techniques, inserting a custom synthesized G-block (IDT) containing a CD28 derived hinge and transmembrane domain linked directly to the CD3ζ intracellular signaling domain into a MSGV1 backbone.

#### Retrovirus generation

Retroviral particles containing the CAR plasmid and RD114 enveloping plasmid were produced by transfecting HEK293-GP cells with Lipofectamine™ 2000 (Invitrogen, cat# 11668027). Supernatant was collected and stored 48- and 72- hours after transfection and used fresh or stored at -80°C.

#### CAR T cell transduction

CAR T cells were produced as previously described[43]. Briefly, leukapheresis cones were purchased from Versiti Blood Center of Wisconsin. T cells were isolated using negative selection with Stem Cell Technologies EasySep™ kit per manufacturer instructions. T cells were either used fresh or cryopreserved with Cryostore™ media (Stemcell Technologies, cat# 11141D) and stored at -150°C. Cryopreserved cells were thawed and activated with aCD3/aCD28 Dynabeads (Thermo Fisher, cat# 11141D) on the same day in AIM V (Thermo Fisher, cat# 12055083) + IL-2 (Thermo Fisher, cat# 200-02). Activated cells were retrovirally transduced with their respective CAR plasmid on days 3 and 4 and

Dynabeads were removed on day 5. Media and IL-2 were changed every 2 to 3 days until day 10, when T cells were frozen in Cryostore$^{TM}$ media.

**Preparation of DNA hairpin-based molecular tension probes on glass surfaces**

Surfaces were prepared with minor modification to previously described protocols[6]. Briefly, no. 1.5H glass coverslips (Ibidi) were sonicated in Nanopure water (3 × 3 min) followed by EtOH (3 × 3 min) and dried at 100 °C for 15 min. They were then etched in piranha solution (3:1 conc. $H_2SO_4$:$H_2O_2$) for 15 min (caution: extremely corrosive) and washed five times with Nanopure water. Slides were then transferred to a beaker containing 3% 3-aminopropyltriethoxysilane (APTES) in ethanol for 1 hour, washed with ethanol thrice, and dried with $N_2$. The slides were then mounted onto a six-channel microfluidic chamber (Sticky-Slide VI 0.4, Ibidi). Into each channel, ~50 µl of NHS-PEG$_4$-azide (10 mg/ml, Click Chemistry Tools) in 0.1 M $NaHCO_3$ (pH 9) was added and left to react for 1 hour. The channels were then washed thrice with 1 ml of Milli-Q water, and then buffer exchanged with 1ml of 1X PBS. The surfaces were blocked with 0.1% BSA in 1X PBS for 30 min and then washed thrice with 1X PBS. Subsequently, the hairpin tension probes were assembled in 1 M NaCl by mixing the hairpin strand (2.2 µM), biotin ligand cy3B strand (2.2 µM), and DBCO BHQ1 strand (2 µM). The mixture was heat-annealed in a thermocycler at 95°C for 5 min and cooled down to 25°C over a 30-min time window. The assembled probe (~50 µl) was added to the channels with ~50 µl of 1X PBS (total volume = ~100 µl) and incubated overnight at RT in dark. This strategy enables covalent anchoring of the tension probes via strain-promoted cycloaddition between the azide functionalized- surface and DBCO on the probe anchor strand. On the following day, unbound probes were washed out with three 1X PBS rinses. Streptavidin (100 µg/mL) was added and incubated for 45 minutes at room temperature, followed by washing thrice with 1X PBS. CAR specific ligand (10 µg/mL) was added, incubated for 45 minutes, and rinsed three times. Before imaging, the surfaces were exchanged into 1X PBS + 2% FBS.

**Image acquisition**

Fluorescence imaging was performed using a Nikon Ti2 Eclipse inverted microscope equipped with a ring total internal reflection fluorescence (TIRF) illumination system (ILAS Ring TIRF). Images were captured using a Quest CMOS camera with an oil immersion, 1.49 NA 100× Apo objective (Nikon Instruments). Imaging parameters were kept constant in the 561 channel with 50% laser power and 200ms exposure time. For MTP imaging, CARs were thawed 30 minutes prior to imaging, stored at 4°C in PBS + 2% FBS and seeded at ~5 x $10^5$ CAR-positive T cells per well 10 minutes prior to image acquisition. For locking experiments, a 1 µM DNA locking strand was added and images were taken immediately following addition of the locking strand and again every 60 seconds for 10 minutes. The 5-minute timepoint was chosen for analysis to avoid signal saturation. For calcium imaging, laser power was reduced to avoid bleaching due to fast acquisition; however, direct comparisons with these fluorescence intensities were not made with any other measurements.

**Image analysis**

For image analysis, the camera dark signal was first subtracted from all images. A flatfield correction was constructed for each experimental replicate by taking images of 4-10 cell free regions, averaging these images, gaussian smoothing the result, and normalizing to the maximum observed intensity to generate the illumination profile of the microscope. Cell tension fluorescence images were then flatfield corrected by dividing by this illumination profile. Regions of interest (ROIs) around cells were then selected and the local background was measured within selected ROIs near the cell. This local background was subtracted from the MTP images. Finally, the images were thresholded using the standard deviation of the cell-free background ROI. Pixels with intensities less than 2 standard deviations above the local, cell-free background fluorescence were set to NaN and removed from further computation. Finally, the remaining pixelwise intensities following background subtraction and thresholding were integrated to produce a single, integrated MTP fluorescence intensity for each cell. This image analysis routine is depicted graphically in **SI Appendix Fig. S12**.

**Cell dynamics analysis**

A custom edge detection algorithm was developed which generated masks from both RICM contact area and tension signal region with fluorescence exceeding 2 standard deviations of background fluorescence intensity. The union of these masks defined the cell boundary, from which centroids were computed. Cell edges were tracked at 100 angular positions per timepoint and smoothed using periodic cubic spline interpolation, with edge contours and centroids overlaid across time to visualize membrane dynamics and cell movement. Angular intensity distributions were quantified in 100 angular bins (3.6° each), with integrated intensity calculated within each angular sector of the combined mask. Results were displayed as polar histogram plots showing the angular distribution of tension signal at each timepoint.

**Fluorescence staining**

Approximately ~5 x $10^5$ CAR-positive T cells were seeded and cultured on the MTP surfaces for 20 minutes in 1xPBS + 2% FBS. The tension was locked for 1 minute using a 1 µM locking strand. Cells were then buffer switched with 1mL of 3.7% paraformaldehyde in 1X PBS and fixed for 15 minutes. The surfaces were gently washed twice with 1X PBS and then permeabilized in 0.1% Triton X-100 (sc-29112, ChemCruz) for 15 minutes followed by two subsequent 1X PBS washes. BSA block of 1% was performed for 60min, followed by two PBS washes. Actin staining was performed per manufacturer recommendations in 1X PBS with 2% BSA at a dilution of 1:400 of Alexa Fluor Plus 647 Actin Phalloidin stain. Surfaces were then washed twice with 1× PBS and imaged. For pZap70 staining, BSA block was performed for 24 hour at 4°C followed by two washes with 1X PBS. Approximately 50 µl of blocking solution was kept inside the channel and 20 µl of A647 Mouse Anti-ZAP70 PY319)/Syk (PY352) stock solution was added to each channel and incubated for 1 hour at RT. Surfaces were then washed thrice with 1× PBS and imaged.

**Calcium imaging**

Frozen CAR T cells were thawed in AIM V + IL-2 media and incubated at 37°C for approximately 3-4 hours. They were centrifuged to remove AIM V and washed with Hank’s Balanced Salt Solution (HBSS, #H9269) twice. A resuspension was made in 1 ml of HBSS containing 0.4% (w/v) pluronic F-127 (#P6867) and 5 µM of Fluo-4 AM (#20551, AAT Bioquest). The mixture was kept at 37°C for 30 minutes in the dark. Afterwards, cells were pelleted by centrifugation at 300 rcf for 10 minutes and resuspended in 1 mL 1xPBS. This was repeated twice and the final suspension was made in 1xPBS containing 1 mM $Ca^{2+}$ and 1 mM $Mg^{2+}$, for a final concentration of 50000 cells in 10 µl. The cells were then kept at 4°C for at least 15 minutes before imaging.

**Pharmacological inhibition studies**

LatB was used at 1µM while Y27632 was used at 40 µM. CAR T cells were seeded on MTP substrates, allowed to spread for 10 min, imaged, and subjected to drug treatment. Drug treated images were acquired after 10min of drug treatment. Ethanol was employed as a vehicle control for LatB treatment.

**Intracellular Cytokine Staining by Flow Cytometry**

Cells were activated with antigen coated Invitrogen Dynabeads™ (cat# 11205D) at a 4:1 bead to cell ratio. Dynabeads™ were washed with PBS and incubated for 10 minutes at room temperature with 0.1 mg of biotinylated antigen (CD19, CD33 or CD123) per million beads. Following incubation, Dynabeads™ were washed with PBS and incubated with CAR T cells at a 4:1 ratio for 4 hours at 37°C, 5% $CO_2$. After bead exposure, cells were prepared for flow cytometry with the Invitrogen FIX & PERM Cell Fixation & Cell Permeabilization Kit per the manufacturer’s instructions. Cells were stained with PE-IFNg (BioLegend, cat# 308703), APC/Vio770- Perforin (Miltenyi Biotec, cat# 130-130-850), BV421-Granzyme (Biolegend, cat# 396413), TNFa-BV605 (Biolegend, cat# 502935), BV711-CD4 (Biolegend, cat# 317440), PE/Cy7-CD8 (BD Biosciences, cat# 566858), AF647-IgG, $F(ab')_2$ (Jackson ImmunoResearch, cat# 109-606-006). Zombie Aqua™ Fixability dye (BioLegend, cat# 423101) was used to exclude dead cells from further analysis. Cells were analyzed with a Cytek Aurora Spectral Flow Cytometer and analyzed in SpectroFlo and FlowJo. For visualization of polyfunctionality, data were imported to SPICE 6 (NIH Vaccine Research Center)[44].

**Incucyte Cytotoxicity Assays**

1 x $10^5$ GFP-positive tumor cells were cocultured with CAR T cells at 1:2, 1:4, and 1:8 effector: target ratios. Cells were cultured in a 96 well plate with 200 mL of complete RPMI media (Sigma Aldrich, cat# R8758) at 37°C for 5 days. 4 images per well were captured every 4 hours with 10x magnification with the Incucyte® S3 Live-Cell Analysis System. Total integrated intensity of green fluorescence intensity was measured, normalized to the starting intensity in each image and plotted over time.

**Data Availability**

Raw data and code used in data analysis are available from the corresponding authors upon reasonable request.

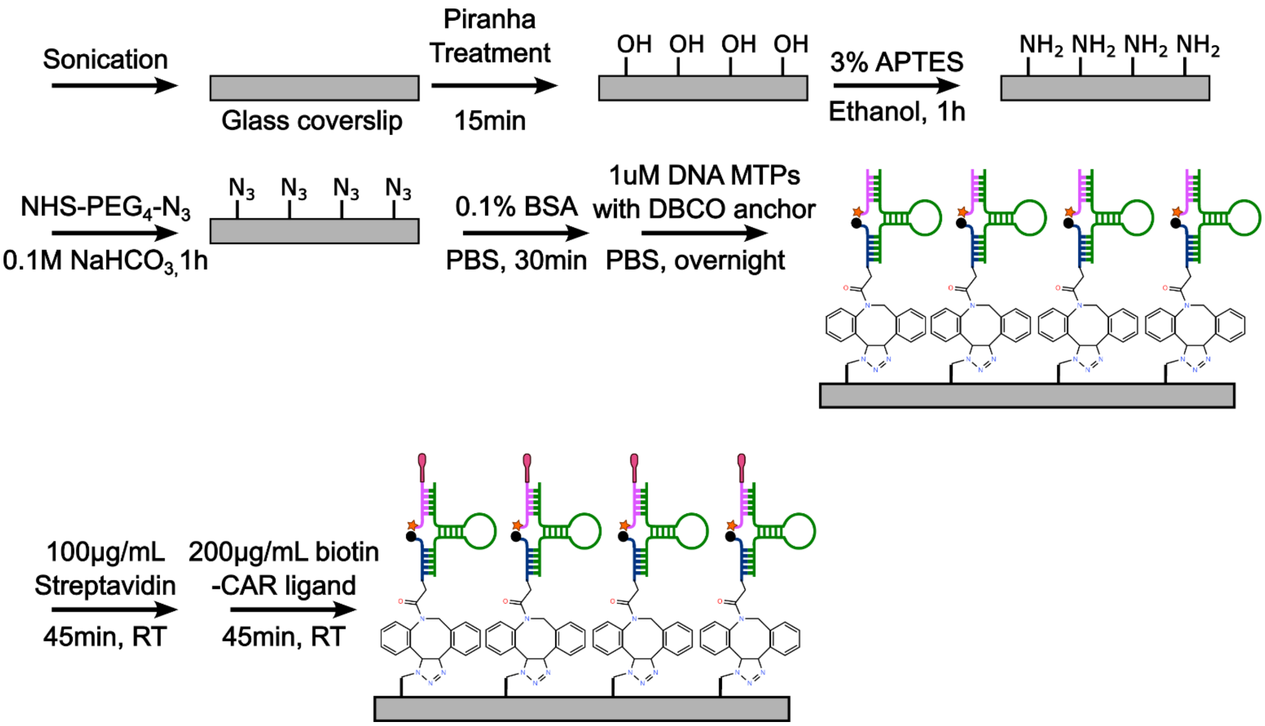


**Figure S1: Surface preparation scheme.** Schematic of the surface functionalization strategy by which DNA MTPs were covalently coupled to glass coverslips and conjugated with biotinylated CAR T cell target protein.

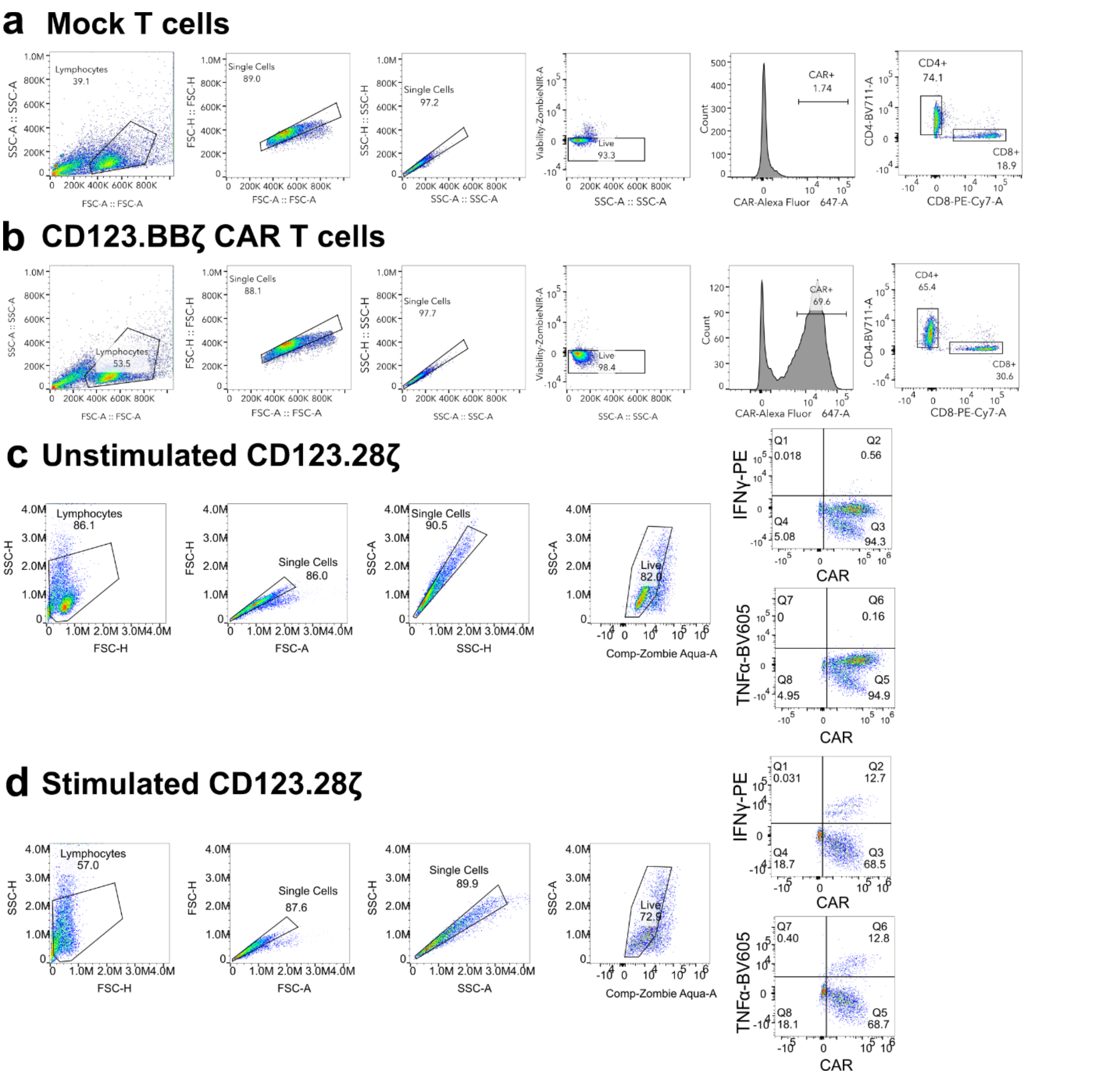


**Figure S2: Flow cytometry gating strategy for CAR T cell phenotyping and intracellular cytokine staining.** Lymphocytes were first gated based on forward scatter area (FSC-A) versus side scatter area (SSC-A), followed by doublet exclusion using FSC-H vs. FSC-A and SSC-H vs. SSC-A to identify single cells. Viable cells were selected based on exclusion of ZombieNIR viability dye. CAR expression was detected using an anti-Fab Alexa Fluor 647-conjugated antibody, and cells were further stained with CD4 (BV711) and CD8 (PE-Cy7) to distinguish T cell subsets. **(a)** Gating strategy for mock T cells, showing background levels of CAR staining. **(b)** Gating strategy for CD123.BBζ CAR T cells, showing a distinct CAR-positive population. Percentage of CAR positive cells is indicated within the live, single lymphocyte population. Gating strategy for intracellular cytokine staining quantification performed in **SI Appendix Fig. S11** for (**c**) unstimulated and (**d**) stimulated with CAR-target ligand coated beads.

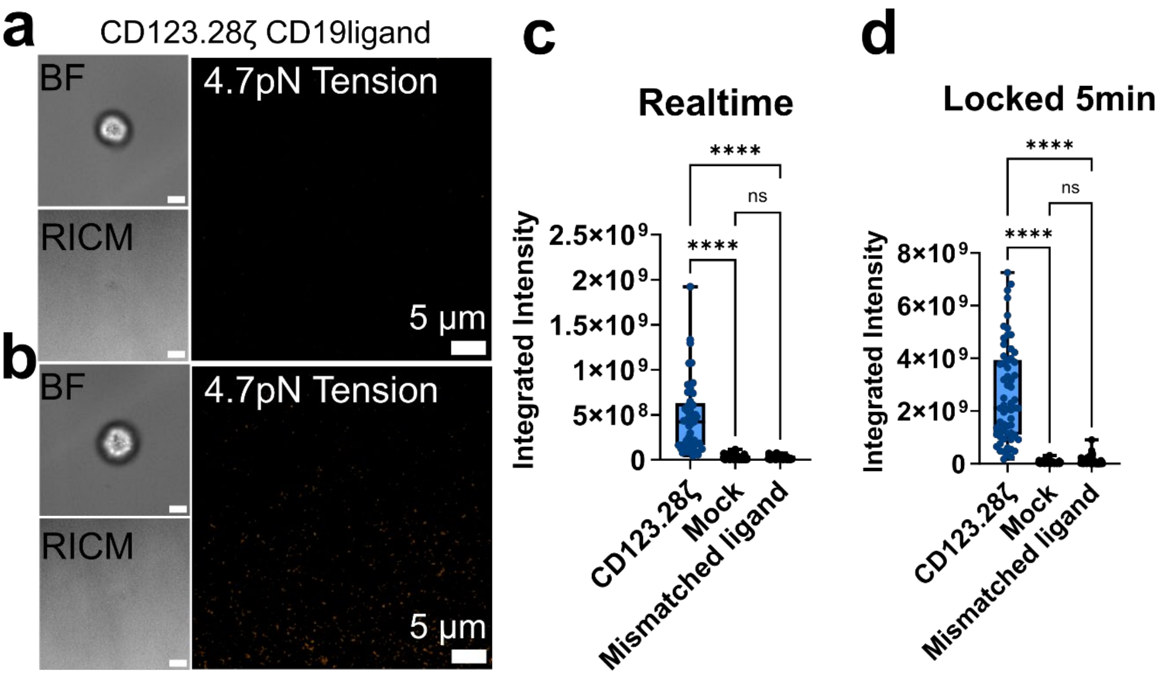


**Figure S3: CARs do not transmit force to irrelevant ligand.** BF, RICM and 4.7pN MTP tension map of a CD123.28ζ CAR T cell on CD19-conjugated DNA-MTPs in (**a**) real time imaging and (**b**) following 5 minutes of signal integration with a DNA "locking strand." Box and whisker plots quantifying the (**c**) real time and (**d**) locked MTP fluorescence signal of CD123.28ζ cells on CD123 conjugated MTPs, mock transduced T cells on CD123 MTPs, and CD123.28ζ CAR T cells on CD19-presenting MTPs (mismatched ligand). Each dot represents the integrated 4.7pN MTP tension fluorescence signal from a single CAR T cell. Data from *n* = 3 independent experiments are aggregated in this plot. Note that CD123.28ζ data in **c** and **d** is replotted from CD123.28ζ data in **Fig. 2e, f** for ease of comparison with mock and mismatched ligand here. Scalebars in BF and RICM in **a** and **b** are 5μm.

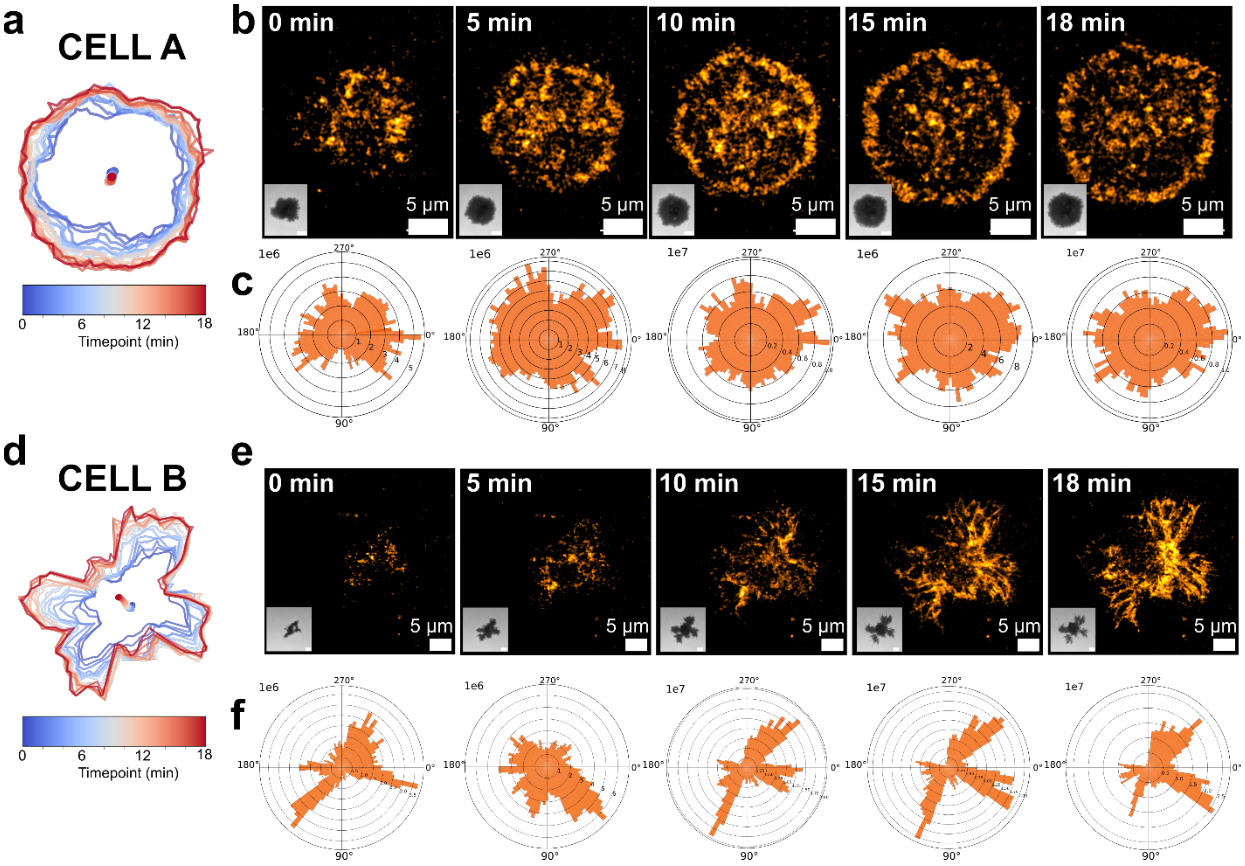


**Figure S4: CARs transmit dynamic forces on MTPs. (a)** Plot depicting the location of the edge of a CD123.28ζ CAR from 0-18 minutes as it spread on a CD123-conjugated MTP surface. The color of the lines indicates the time at which the cell occupied a given location. The colored dots in the middle of the cell indicate the centroid position as a function of time. **(b)** Thresholded 4.7pN MTP fluorescence images of the same cell shown in **a**. The inset images shows cell-surface contact area via RICM. (**c**) Angular histogram of the distribution of MTP fluorescence. **(d-f)** Another cell (Cell B) subject to the same analysis. This analysis is representative of ~20 cells from *n* = 3 independent experiments. Scalebars in RICM insets in **b** and **e** are 5μm.

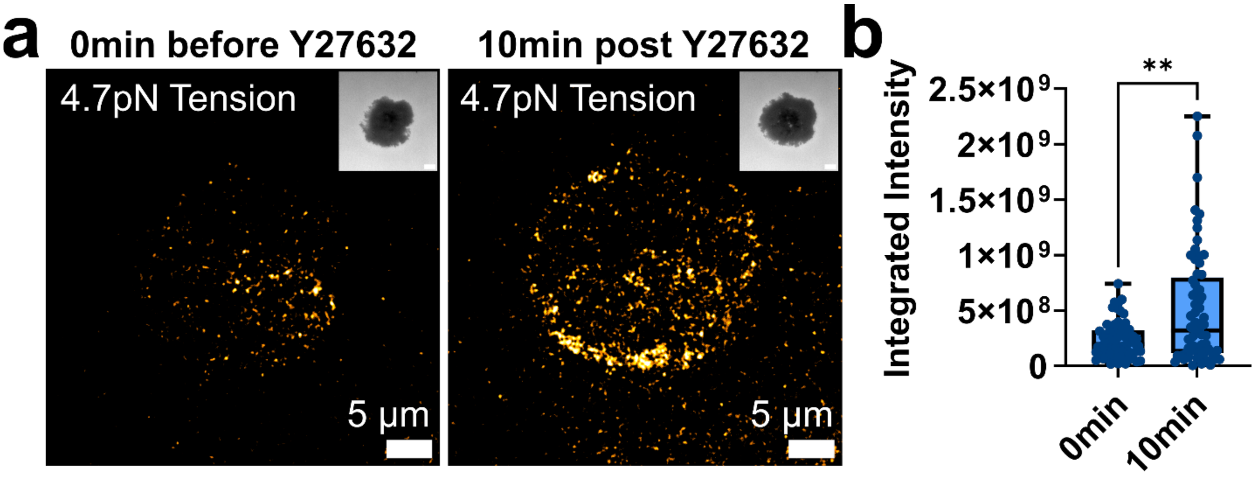


**Figure S5: Forces exerted by CARs are not myosin-mediated. (a)** 4.7pN MTP fluorescence image of a CD123.28ζ CAR T cell at T=0 min post addition and T=10 min post Y27632 drug addition, RICM as inset. **(b)** Quantification of MTP fluorescence before and after addition of drug. Images in **a** are representative of *n* = 3 independent experiments conducted with CARs generated from 1 human donor. **** $p<0.0001$, Mann-Whitney test. Scalebars in RICM insets in **a** are 5μm.

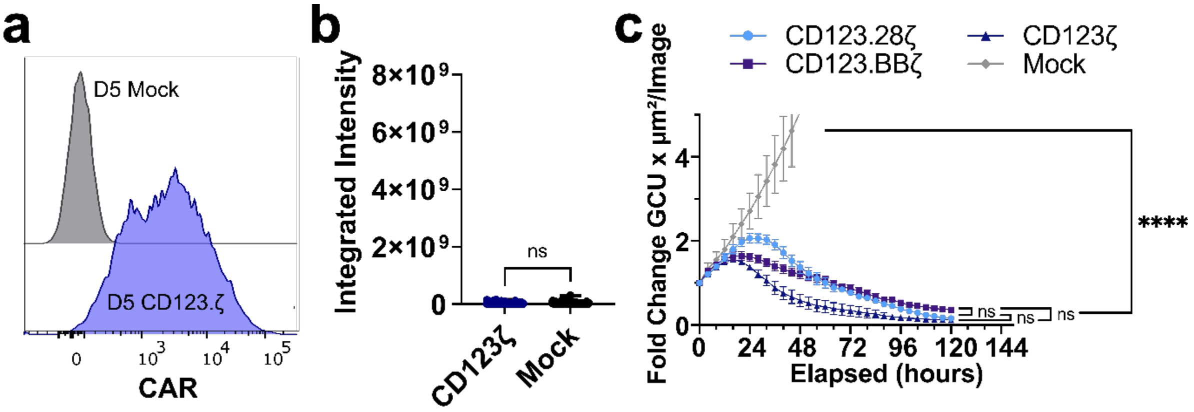


**Figure S6: "First-generation" CD123.ζ CARs fail to generate forces exceeding 4.7pN but retain cytotoxicity**

(**a**) Flow cytometry quantification of CAR expression on live, single cells in CD123.ζ first-generation CAR and mock transduced T cells. (**b**) Box and whisker plots in which each dot represents the integrated 4.7pN MTP tension fluorescence signal from CD123.ζ CAR T cell and mock transduced T cells. (**c**) Fold change in per well GFP fluorescence at t=0 measured over 120 hours in a single donor across *n=3* replicate wells of CD123.28ζ, CD123.BBζ, and CD123ζ CAR T cells reveal no difference in their cytotoxicity. Data in **a** is from one representative donor. ns = not significant, * = p<0.0001. Statistical significance was assessed via Mann-Whitney test and Kruskal-Wallis test.

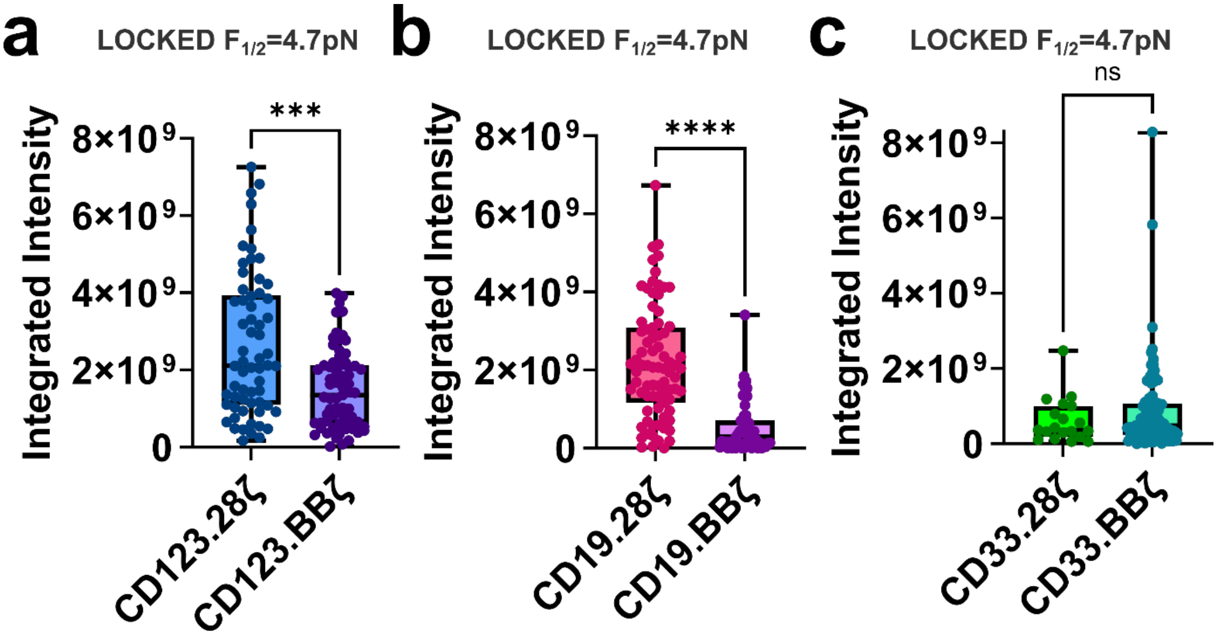


**Figure S7: Both CD28 and 4-1BB co-stimulatory domain CAR T cells targeting CD123, CD33, or CD19 transmit pN forces to target ligand.** Box and whisker plots quantifying the whole cell integrated MTP fluorescence signal following 5 minutes of locking with $F_{1/2}$=4.7pN for (**a**) CD123.28ζ versus CD123.BBζ (n=3 donors) (**b**) CD19.28ζ versus CD19.BBζ, (n=2 donors) (**c**) CD33.28ζ versus CD33.BBζ, (n=2 donors). ns = not significant, *** = $p<0.001$, **** = $p<0.0001$.

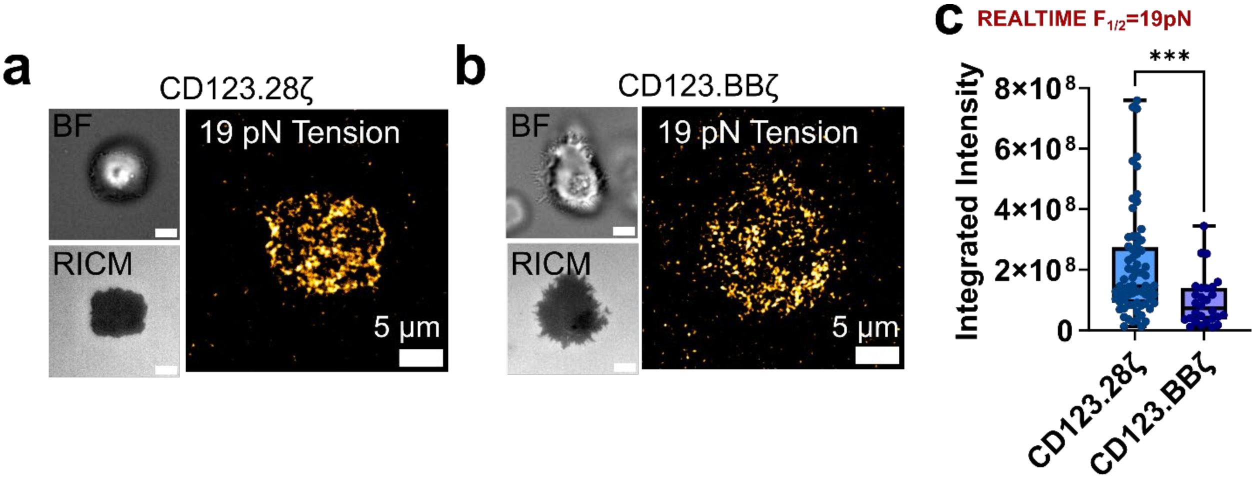


**Figure S8: CD123 CAR T cell exert forces exceeding 19pN.** BF, RICM, and 19pN MTP fluorescence image of (**a**) CD123.28ζ and (**b**) CD123.BBζ CAR T cells seeded on CD123-presenting MTPs. (**c**) Quantification of realtime MTP fluorescence with $F_{1/2}$=19pN for CD123.28ζ versus CD123.BBζ. ***=$p<0.001$. Scalebars in BF and RICM in **a** and **b** are 5μm.

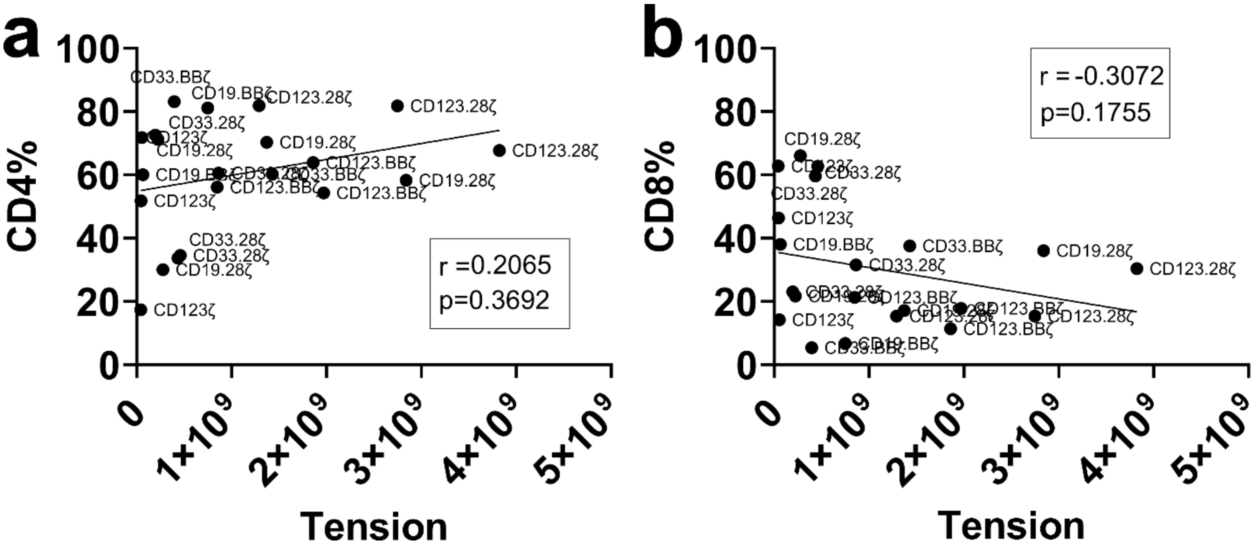


**Figure S9: MTP tension fluorescence comparison with CD4/8 ratios of CAR T cells.** Flow cytometry quantification of CAR-positive cell phenotype versus average integrated MTP fluorescence per CAR product. Tension in these plots indicates the average integrated MTP fluorescence for all measured cells from each CAR T cell product. No statistical correlation was measured between average integrated MTP fluorescence and (**a**) %CD4$^+$ (**b**) %CD8$^+$.

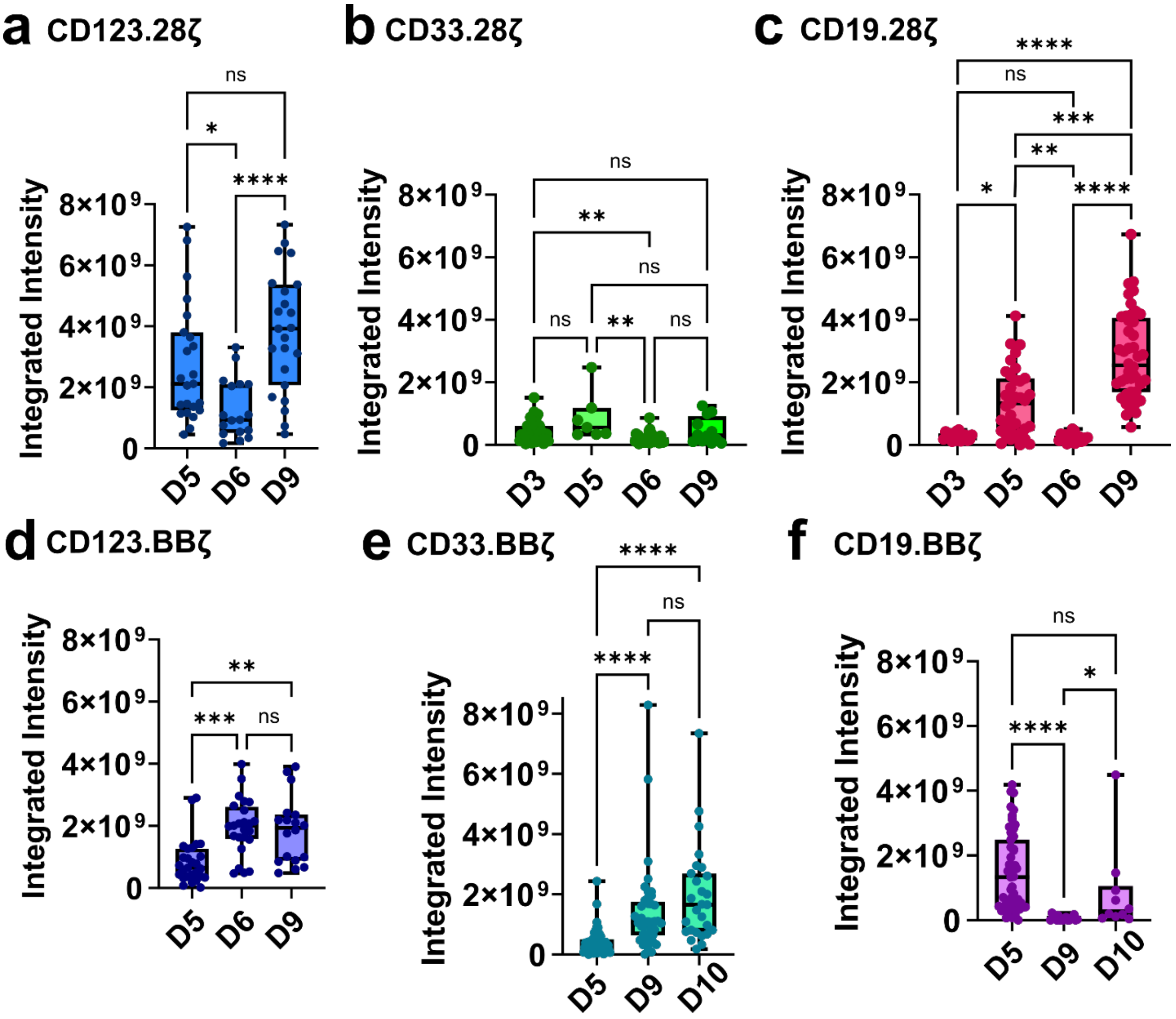


**Figure S10: Tension of CARs is donor T cell-dependent.** (**a-f**) Box and whisker plots in which each dot represents the integrated 4.7pN MTP tension fluorescence signal from a single CAR T cell following 5 minutes of locking. The nature of the CAR product is denoted by the following naming convention: target protein (e.g. CD123), co-stimulatory domain (if present, 28 for CD28 and BB for 4-1BB), ζ for the CD3 zeta domain. The x axis of each plot reflects de-identified donor number as the letter D followed by a number. ns = not significant, * = $p<0.05$, ** = $p<0.01$, *** = $p<0.001$, **** = $p<0.0001$. Statistical significance was assessed via the Kruskal-Wallis test.

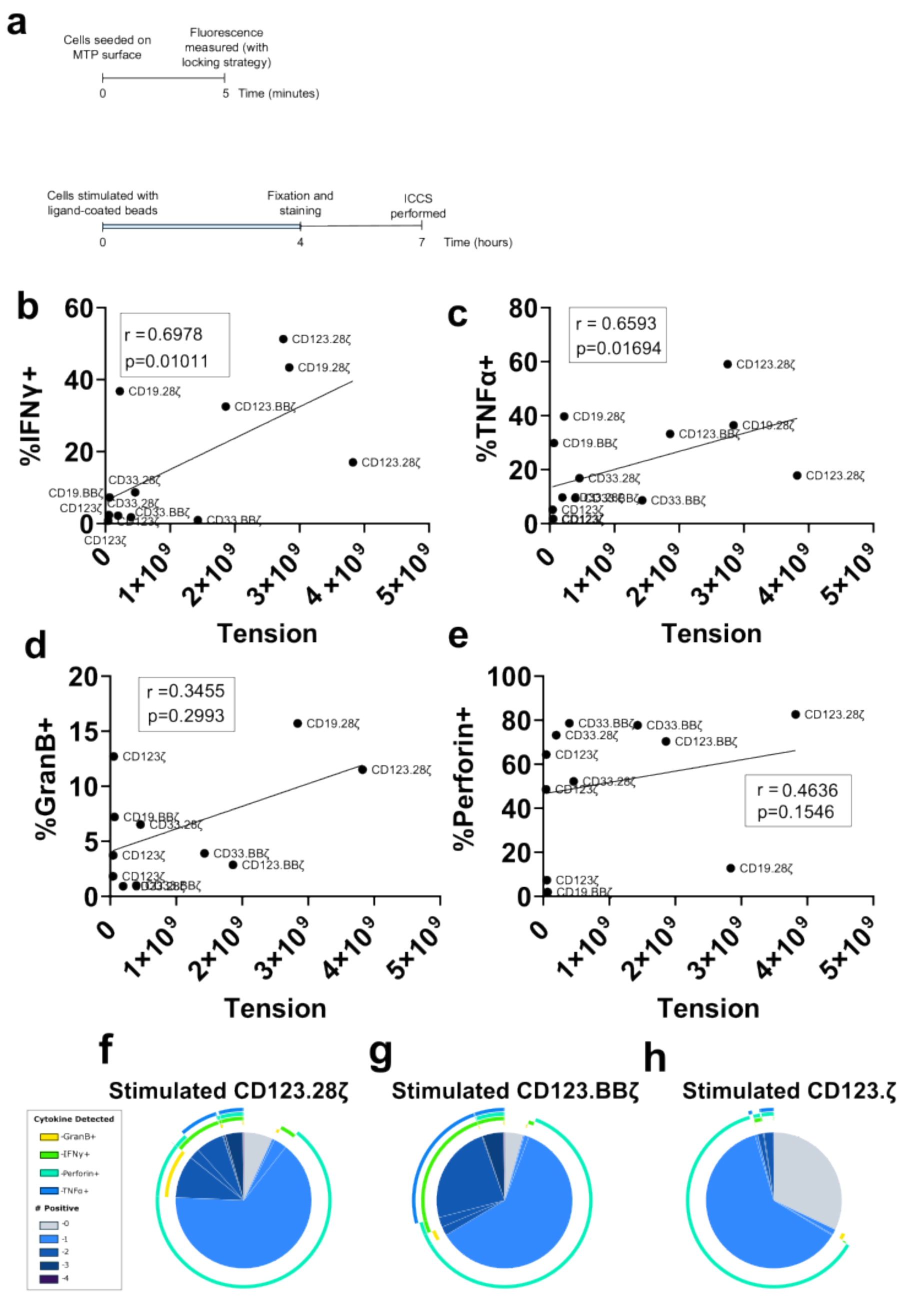


**Figure S11: Intracellular cytokine staining following CAR target ligand-conjugated bead stimulation. (a)** Experimental timeline in which the same CAR product (defined as a unique combination of CAR construct and donor) were subjected to both MTP fluorescence imaging and intracellular cytokine staining (ICCS) for IFNγ, TNFα, GranB, and Perforin following stimulation with CAR target-ligand conjugated beads. (**b-e**) Plot of mean integrated (locked 5min) intensity for each CAR product (x-axis) plotted against (**b**) %IFNγ+, (**c**) %TNFα+, (**d**) %GranB+, and (**e**) %Perforin+ CAR T cells for each CAR product following *in vitro* bead stimulation. CAR product identity for each point is provided on the plot. Boolean analysis of (**f**) CD123.28ζ, (**g**) CD123.BBζ, and (**h**) CD123.ζ ICCS of representative live, CAR positive cells.

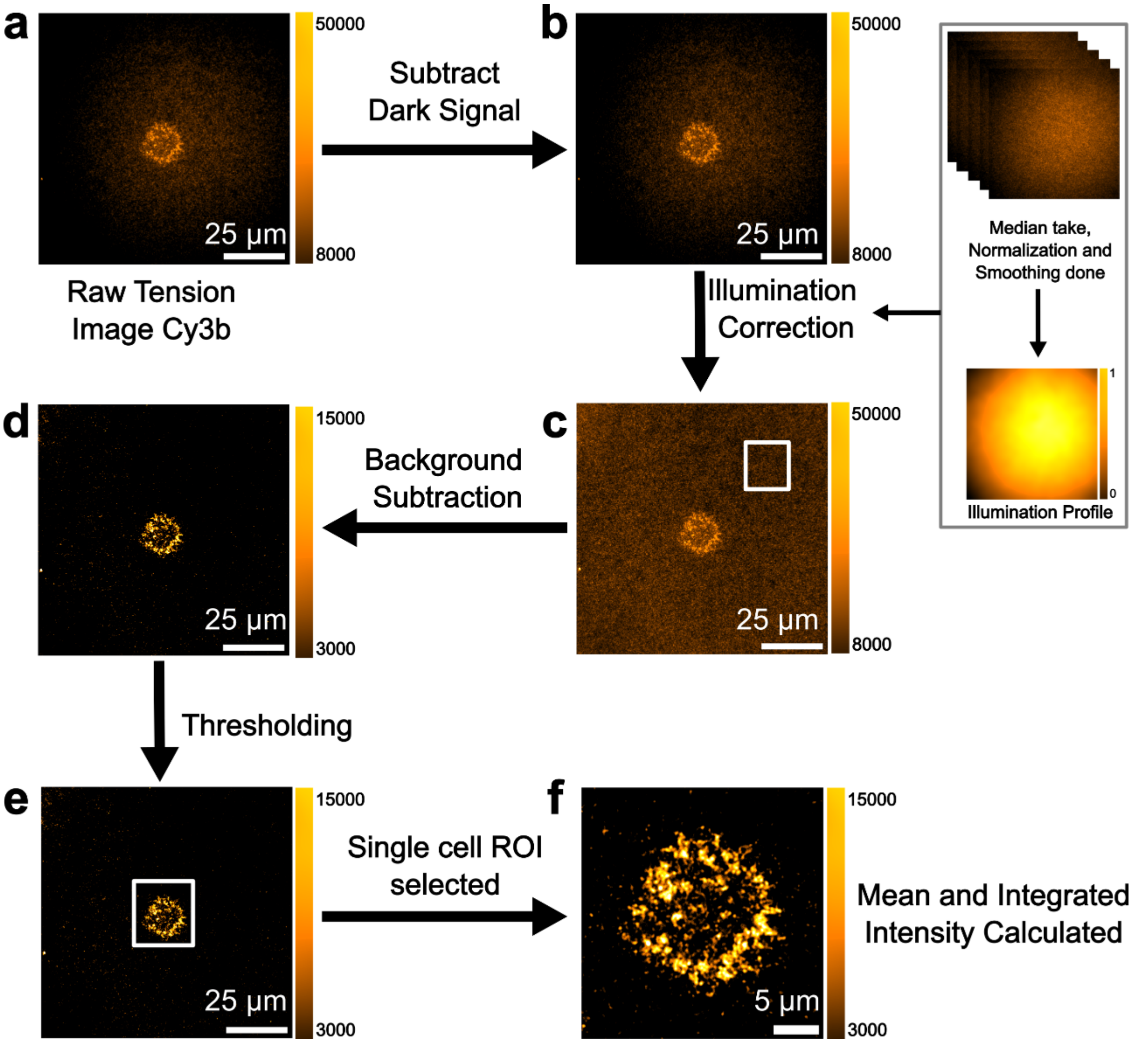


**Figure S12. Image processing scheme.** (**a**) Raw Cy3b-labeled molecular tension probe (MTP) image of a single CAR T cell. (**b**) Dark signal subtraction applied to remove camera baseline noise. (**c**) Illumination correction using a normalized, smoothed illumination profile generated from median projections of blank field images. (**d**) Local background subtraction performed using an off-cell ROI. (**e**) Thresholding was applied to isolate the cell region of interest (ROI). The threshold was set as 2 times the standard deviation of the cell-free background. Thresholded pixels were set to NaN for downstream analysis. (**f**) Final ROI used for quantification, with mean and integrated fluorescence intensities calculated.

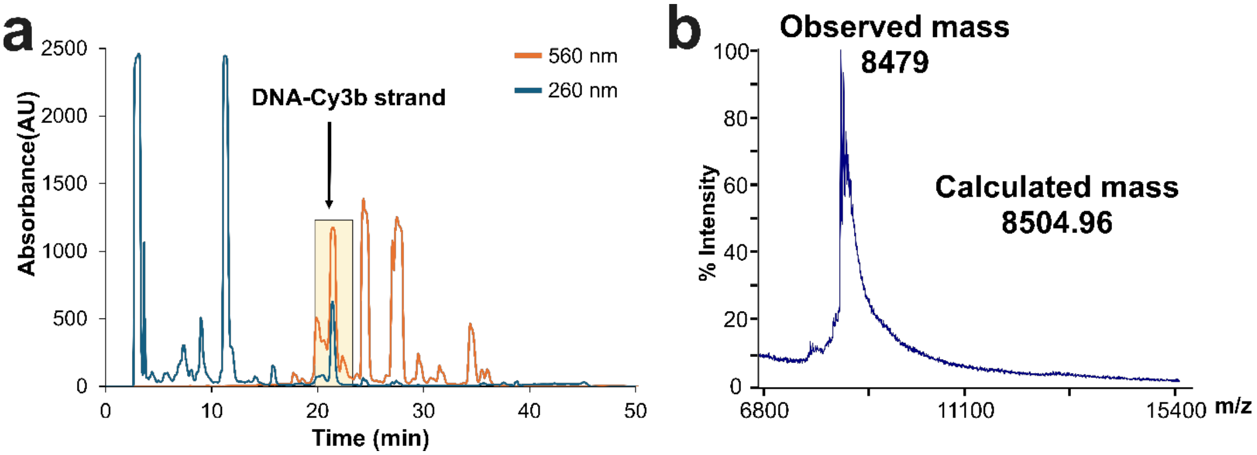


**Figure S13. Dye labelled DNA strand synthesis.** (**a**) Representative HPLC trace and (**b**) MALDI-TOF spectrum of Cy3b labelled biotin ligand DNA strand.

**Table S1. List of oligonucleotides**

| Sequence ID | Description | Sequence (5' to 3') |
|---|---|---|
| 22GC hairpin | 4.7pN hairpin strand in MTPs | GTG AAA TAC CGC ACA GAT GCG TTT GTA TAA ATG TTT TTT TCA TTT ATA CTTTAA GAG CGC CAC GTA GCC CAG C |
| biotin ligand amine | ligand strand on top | /5Biosg/TT TGC TGG GCT ACG TGG CGC TCT T/3AmMO/ |
| biotin ligand cy3B | Cy3B labeled ligand strand on top | /5Biosg/TT TGC TGG GCT ACG TGG CGC TCT T - Cy3B |
| DBCO BHQ1 strand | bottom and quencher strand with DBCO | /5BHQ_1/CGC ATC TGT GCG GTA TTT CAC TTT /3DBCON/ |
| 100GC hairpin | 19pN hairpin strand in MTPs | GTG AAA TAC CGC ACA GAT GCG TTT GCG CGC GCG CGC TTT TGC GCG CGC GCG CTT TAA GAG CGC CAC GTA GCC CAG C |
| locking strand | complement to partial hairpin | GAA AAA AAC ATT TAT ACC CTA CCT A |

**Table S2. List of fluorophores and antibodies utilized in flow cytometry**

| Target | Fluorophore | Manufacturer | Catalog # | Lot # |
|---|---|---|---|---|
| PD-1 | BV605 | Biolegend | 367426 | B422619 |
| TIM-3 | BV421 | Biolegend | 345008 | B456511 |
| LAG-3 | PE | Biolegend | 369306 | B387558 |
| CD4 | BV711 | Biolegend | 317440 | B456296 |
| CD8 | PE/Cyanine 7 | Biolegend | 344750 | B426650 |
| IFNg | PE | Biolegend | 502508 | B449171 |
| Perforin | APC-Vio770 | Miltenyi Biotec | 130-130-850 | 1324030941 |
| TNFa | BV605 | Biolegend | 502935 | B406341 |
| Granzyme B | BV421 | Biolegend | 396413 | B407094 |
| Viability | ZombieNIR | Biolegend | 423105 | B440686 |
| Viability | Zombie aqua | Biolegend | 77143 | B451565 |
| IgG, $F(ab')_2$ fragment specific | Alexa Fluor® 647 | Jackson ImmunoResearch | 115-606-072 | 163053 |
